# Hippocampus plays a role in speech feedback processing

**DOI:** 10.1101/2020.03.21.001792

**Authors:** Vincent van de Ven, Lourens Waldorp, Ingrid Christoffels

**Affiliations:** Department of Cognitive Neuroscience, Faculty of Psychology and Neuroscience, Maastricht University, P.O. Box 616, 6200 MD, Maastricht, The Netherlands; University of Amsterdam, Amsterdam, The Netherlands

**Keywords:** Overt picture naming, hippocampus, expectation, speech monitoring

## Abstract

There is increasing evidence that the hippocampus is involved in language production and verbal communication, although little is known about its possible role. According to one view, hippocampus contributes semantic memory to spoken language. Alternatively, hippocampus is involved in the processing the (mis)match between expected sensory consequences of speaking and the perceived speech feedback. In the current study, we re-analysed functional magnetic resonance (fMRI) data of two overt picture-naming studies to test whether hippocampus is involved in speech production and, if so, whether the results can distinguish between a “pure memory” versus an “expectation” account of hippocampal involvement. In both studies, participants overtly named pictures during scanning while hearing their own speech feedback unimpededly or impaired by a superimposed noise mask. Results showed decreased hippocampal activity when speech feedback was impaired, compared to when feedback was unimpeded. Further, we found increased functional coupling between auditory cortex and hippocampus during unimpeded speech feedback, compared to impaired feedback. Finally, we found significant functional coupling between a hippocampal/supplementary motor area (SMA) interaction term and auditory cortex, anterior cingulate cortex and cerebellum during overt picture naming, but not during listening to one’s own pre-recorded voice. These findings indicate that hippocampus plays a role in speech production that is in accordance with an “expectation” view of hippocampal functioning.

## Introduction

Traditionally, language has been viewed as a fronto-temporal neocortical function and studied in isolation from memory. However, there is growing evidence that the medial temporal lobe (MTL), particularly the hippocampus, also contributes to speech production and verbal communication (Duff et al., 2008; Duff and Brown-Schmidt, 2012; Covington and Duff, 2016; Llorens et al., 2016; Piai et al., 2016; Kepinska et al., 2018). One interpretation of this evidence is that verbal communication bears on declarative memory processes that are governed by the hippocampus (Squire and Zola, 1998; Manns et al., 2003; Klooster and Duff, 2015). Alternatively, the hippocampus supports control of speech production by generating context-based expectations of future perceptual events or actions (Friston and Buzsáki, 2016; Stachenfeld et al., 2017), which can then be matched to speech feedback. One scenario in which these two views of the hippocampus in language processing can be separated is when the speech monitoring process during speaking is impaired. When we speak, we continuously monitor our speech production by comparing what we intended to say to our own speech output (speech feedback) (Levelt, 1983; Indefrey and Levelt, 2000; Schiller and de Ruiter, 2004). A mismatch between actual and expected sensory (auditory) consequences of speech acts can result in more cognitive control to optimize future speech acts. Such a mismatch can result from internal causes, such as (incidental) poor organization or control of motor functions while stuttering, or external causes, in which loud environmental noise masks or alters speech feedback. Adopting a “pure memory” view of hippocampal function one would predict no alternation of hippocampal activity with mismatching speech feedback, as the hippocampus contributed semantic or associative memory content to the speech formation but is not involved in monitoring speech consequences. Other brain areas may monitor how well the speech production matches the sensory consequences of that output (e.g., (Tourville et al., 2008; Hickok, 2012)). From the perspective of the “expectation” view of hippocampal function, impaired speech feedback will alter hippocampal activity, as the hippocampus is actively involved in processing the degree of mismatch between perceived feedback to sensory expectations derived from memory (Kumaran and Maguire, 2009; de Lange et al., 2018). In the current study, we tested the involvement of the hippocampus in speech monitoring by analysing whether hippocampal activity and connectivity with areas of the speech monitoring network changes with changing speech feedback quality.

Studies that investigated the neural correlates of speech monitoring during speech production showed increased activity in auditory cortex and superior temporal gyri, lateral and medial prefrontal cortex and premotor areas, when speech feedback was impaired or interrupted, compared to unimpeded speech feedback (McGuire et al., 1996; Hirano et al., 1997; Hashimoto and Sakai, 2003; Heinks-Maldonado et al., 2005; Christoffels et al., 2007, 2011; Tourville et al., 2008; Zheng et al., 2010). This effect is not observed when participants listen to their own pre-recorded voice (Zheng et al., 2010; Christoffels et al., 2011). These findings imply that the motor system that plans and coordinates speech acts provides a prediction of the sensory consequences of those speech acts (Hickok, 2012; Skipper et al., 2017), which is then compared to sensory feedback representations in auditory cortex. Indeed, several components of the (sub)cortical motor system have been implicated in providing such a predictive feed-forward signal for sensory comparison in speaking as well as other self-initiated actions (Wolpert et al., 1995; Blakemore et al., 1998; Heinks-Maldonado et al., 2005; McNamee and Wolpert, 2019). Particularly, the supplementary motor area (SMA) may be important in this process. Neurophysiological recordings showed that SMA neurons can code for future movements when they are part of a learned sequence of movements (Tanji and Shima, 1994). Further, SMA activity correlates with the voluntary generation of covert auditory speech (speech imagery) (Lima et al., 2016) as well as the involuntary generation of auditory verbal hallucinations of voices in schizophrenia (van de Ven, 2012). Human clinical studies showed that traumatic or pathological lesions of the SMA resulted in diminished speech control (Jonas, 1981; Alario et al., 2006; Hertrich et al., 2016) while transcranial magnetic stimulation (TMS) of (pre-)SMA resulted in decreased word production or control of oral gestures (Tremblay and Gracco, 2009), as well as impaired monitoring of self-generated actions (Haggard and Whitford, 2004; Moore et al., 2010). Several fMRI studies showed increased SMA activity (and other frontal and temporal areas) when healthy participants made speech production errors, compared to correct speech acts (Abel et al., 2009; Gauvin et al., 2016). Finally, functional connectivity between SMA and speech perception areas is increased when sensory feedback during speaking is masked, compared to when it is unimpeded (van de Ven et al., 2009), but decreased when articulation errors in degenerative speech disorders increase (Botha et al., 2018), which provides further evidence speech monitoring and control encompass a neural interaction between sensory and motor systems.

If the hippocampus contributes to speech monitoring, then it can be expected that hippocampal activity or connectivity with auditory-motor areas involved in speech monitoring changes when speech feedback is impaired. To test this hypothesis, we reanalyzed brain activity from two previously published speech monitoring studies in which speech feedback was masked by varying levels of superimposed acoustic noise. Importantly, both studies contained the same conditions of noise-masked and noise-free speech feedback, which allowed the data to be aggregated and analysed with the same statistical contrast. A further benefit was that by combining the datasets we doubled sample size and consequently the statistical power. We analysed the combined dataset in three ways. First, we investigated whether hippocampal activity changed when speech feedback was masked, compared to when feedback was unimpeded. A change in hippocampal activity when speech feedback conditions changed can be considered as evidence for the “expectation” view of hippocampal functioning in speech production. Second, we investigated whether hippocampus was functionally connected to the auditory cortex and auditory-motor interaction during speech feedback conditions. Task-dependent functional coupling between auditory cortex and hippocampus would be evidence that hippocampus contributes to the mismatch processing in auditory cortex during speech feedback. Finally, we explored the whole-brain functional network that was associated to the auditory-motor interaction, in order to investigate which areas would be sensitive to an associative mismatch signal that would be represented by the functional coupling between SMA and hippocampus.

## Methods

For this study, we re-analysed two fMRI datasets, Study 1 ((Christoffels et al., 2007), N=12) and Study 2 ((Christoffels et al., 2011), N=11), in both of which participants completed a similar speech monitoring task. Data of both studies were collected using comparable scanning parameters at equal magnetic field strengths (see Table 1). The aggregated data sample contained 23 participants, all of whom provided informed consent prior to the start of the respective experiment. Local ethics committees of Nijmegen University Medical Center (Study 1) or the Faculty of Psychology and Neuroscience (Study 2) approved the studies.

**Table 1.**
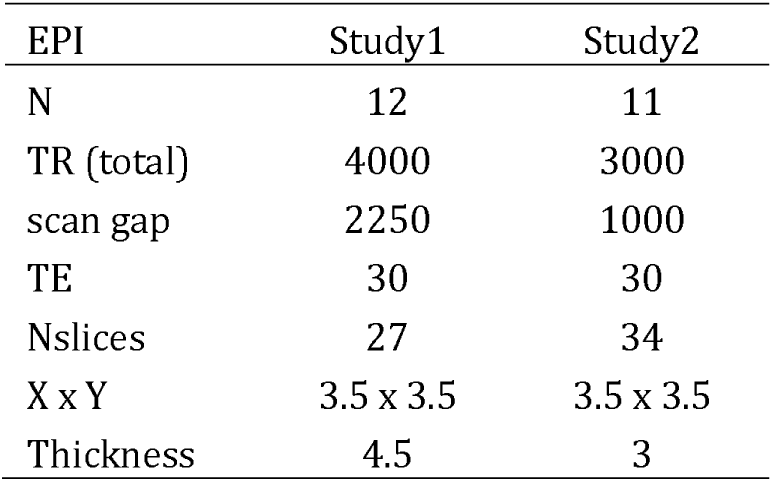
EPI parameters for Study 1 and Study 2. Temporal parameters (TR, TE, scan gap) are listed in milliseconds. In-plane resolution (X x Y) and slice thickness are listed in millimeters.

The experimental designs of both studies were comparable. Participants had to overtly pronounce the name of an object that was shown as a black-and-white line drawing (picture-naming conditions, PN), or heard their own pre-recorded voice speaking out the name of the presented object (listening conditions, LIS). Figure 1 provides a schematic overview of the task design for both studies. During the experiment, acoustic noise was superimposed during either overt picture naming or listening at various volume levels (dB). In Study 1, the acoustic noise volume was either 0 dB or at an individual-tailored volume at which participants could not hear their own speech feedback with overtly speaking (max dB). In Study 2, acoustic noise volumes varied parametrically between 0 and max dB across four levels during picture naming (PN0, PN1, PN2 and PNMax) or listening (LIS0, LIS2, LIS3 and LISMax), which was again individually tailored. A further difference between the studies was the presentation of a covert naming condition in Study 1, which was entirely absent from Study 2. This condition was modelled in the analyses but always excluded from statistical contrasts (see below).

**Figure 1.**
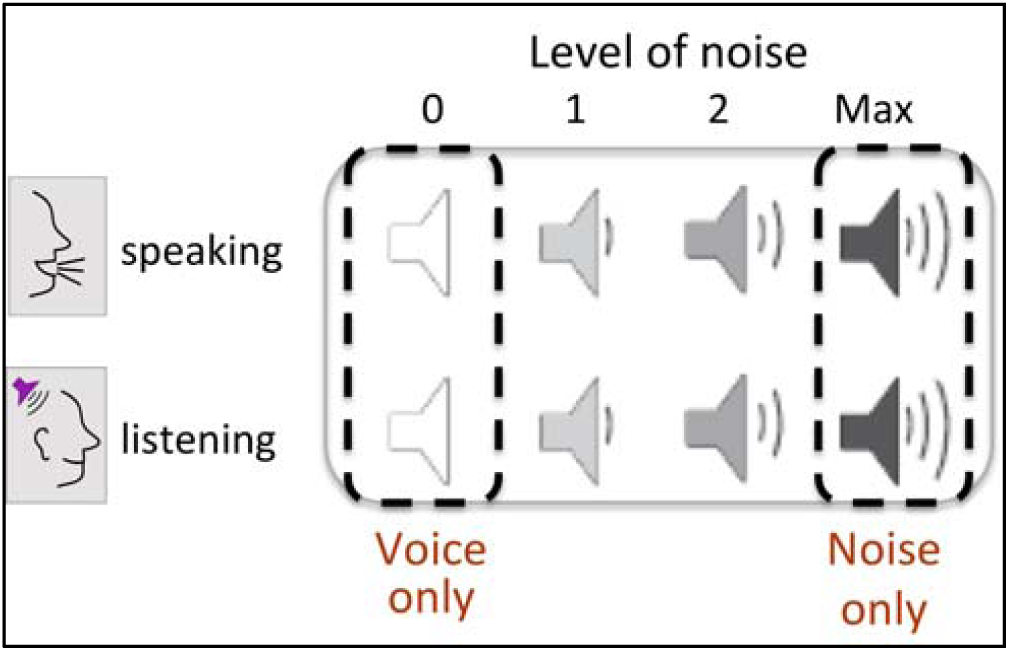
Picture naming conditions used for analysis. In both studies, participants saw pictures and either overtly named them (speaking conditions) or listened to their own pre-recorded naming of those pictures (listening conditions) while acoustic noise was superimposed at various volume levels. In Study 1, noise levels varied between 0 (no superimposed noise) and maximum level (black boxes), while in Study 2 noise levels varied parametrically across four levels (grey box). For the analysis, we chose the Voice only and Noise only conditions that overlapped in both studies.

### Imaging parameters

MR images of both studies were collected on a 3T scanner from the same manufacturer (Siemens Medical Systems; Study 1: Magnetom Trio whole-body at Nijmegen, Study 2: Magnetom Allegra head-only at Maastricht). In both studies, functional volumes were acquired using a T2*-weighted echoplanar imaging (EPI) sequence optimized for blood oxygenation level-dependent (BOLD) contrast (see Table 1 for parameter values of each study). The TR included a silent gap (sparse sampling method (Hall et al., 1999); see Table 1) to allow participants to overtly pronounce the name of the visually presented object without interference of scanner noise during EPI acquisition. High-resolution (1 × 1 × 1 mm^3^) anatomical scans were acquired using a T1-weighted 3D MP-RAGE (Study 1: 192 sagittal slices, TR=2.3s, TE=3.93ms) or an MDEFT sequence (Study 2: 192 sagittal slices, TR= 7.92s, TE=2.4ms).

### Image analysis

All anatomical and functional images were preprocessed using BrainVoyager QX (Goebel et al., 2006) and the NeuroElf toolbox (www.neuroelf.net) and custom-written scripts in Matlab (www.mathworks.com). Preprocessing functional images included slice scan time correction, three-dimensional (3D) head-movement assessment and correction using rigid body transformations, linear trend removal and temporal high-pass filtering (cutoff at appr. 0.004 Hz). The estimated translation and rotation parameters for head-movement were inspected and never exceeded 3 mm or degree. No spatial smoothing was applied. Preprocessed functional timeseries were coregistered to the within-session anatomical 3D dataset using position parameters from the scanner and manual adjustment, and were subsequently spatially normalized to Talairach space (Talairach and Tournoux, 1988) at an iso-voxel resolution of 3 × 3 × 3 mm^3^.

The functional data were then further cleaned using a temporal principal component analysis-based correction procedure (CompCor; (Behzadi et al., 2007)). For each participant, we first estimated temporal noise sources (based on temporal signal-to-noise estimates) from the functional timeseries using pre-defined anatomical templates of the ventricles and white matter, as obtained by tissue segmentation procedures. The temporal noise sources were then subsequently removed from the functional imaging data using a least-squares solution. The normalized and cleaned data were then analysed using a region-of-interest (ROI) approach.

### Regions of interest (ROIs)

ROIs were defined for bilateral auditory cortex (AC), bilateral SMA and bilateral hippocampus in the following ways. To optimize localization of the speech monitoring effect in AC, we used the empirical result of the difference between PN0 and PNMax conditions of Study 1. We note that this selection does not pose a non-independence threat to our current analysis, because 1) this ROI had not been used previously for the analysis of Study 2, and 2) the main target of the current study was to analyse hippocampal activity and functional connectivity. For hippocampal and SMA ROIs, we used the respective maps of the probability atlas by Eickhoff and colleagues (Eickhoff et al., 2005). Specifically, we defined SMA by taking two spheres of 6 mm radius at the three-dimensional center of the left and right Brodmann Area 6 (BA6) (Geyer, 2004). We defined left and right hippocampus by thresholding the probability maps at 60% (Amunts et al., 2005). The SMA and hippocampus maps were aligned to the Talairach spatial template using a 12-parameter affine transformation. The ensuing ROIs superimposed on an average template of all participants are shown as insets in Figure 2, and details are listed in Table 2.

**Table 2.**
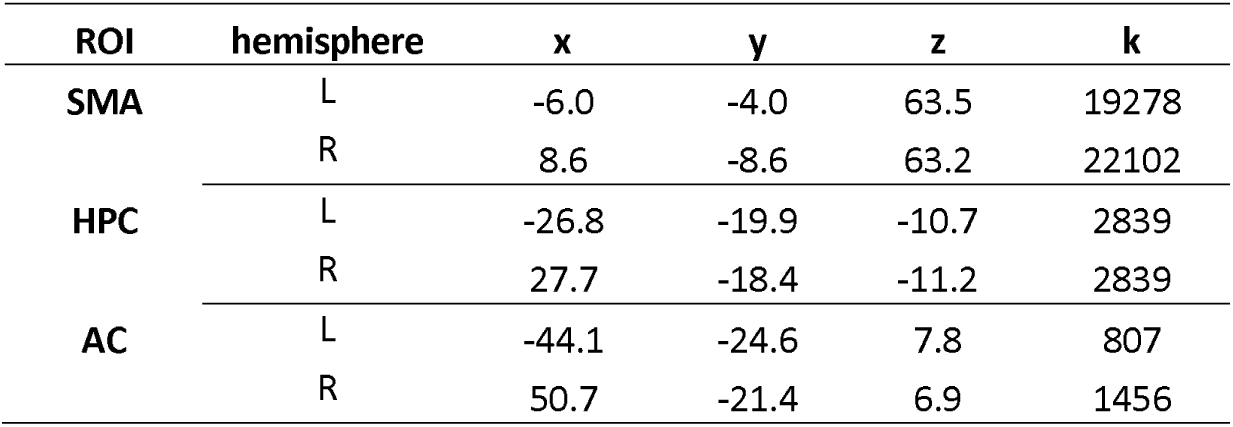
ROI details. ROI coordinates are listed in mm MNI space, ROI size (k) is listed in mm^3^.

| ROI | hemisphere | x | y | z | k |
| --- | --- | --- | --- | --- | --- |
| SMA | L | -6.0 | -4.0 | 63.5 | 19278 |
|  | R | 8.6 | -8.6 | 63.2 | 22102 |
| HPC | L | -26.8 | -19.9 | -10.7 | 2839 |
|  | R | 27.7 | -18.4 | -11.2 | 2839 |
| AC | L | -44.1 | -24.6 | 7.8 | 807 |
|  | R | 50.7 | -21.4 | 6.9 | 1456 |

**Figure 2.**
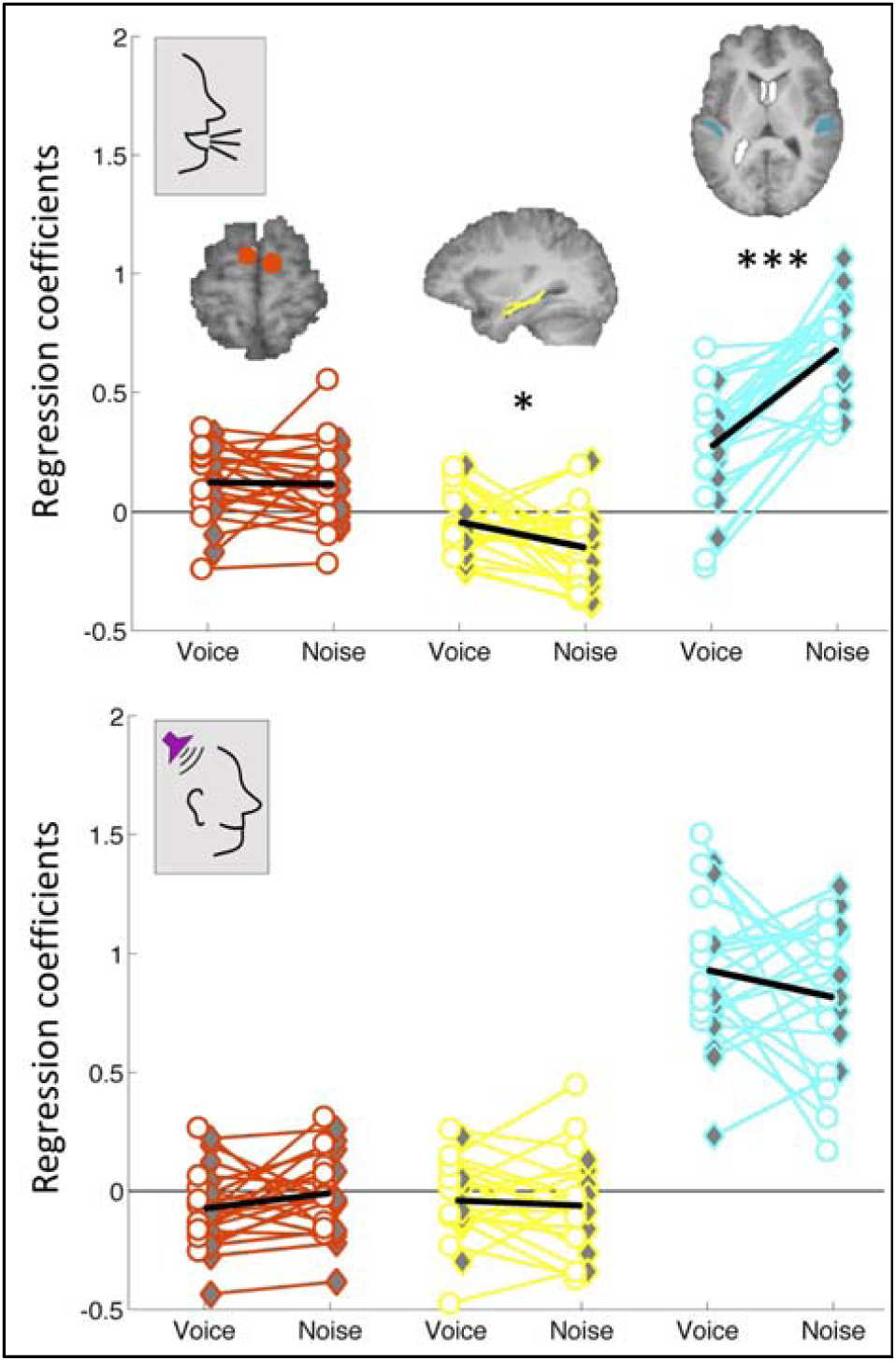
ROI results. Regression coefficients for each participant are shown for supplementary motor area (SMA; in red), hippocampus (yellow) and auditory cortex (blue) during picture naming (upper panel) or listening conditions (lower panel) without or with superimposed acoustic noise (resp., Voice and Noise). ROIs are shown on anatomical brain slices as insets. Study 1 data are marked in grey diamonds, Study 2 data are marked in white circles. * P < 0.05, *** P < 0.001.

### Functional data analysis

Functional timeseries were sampled from and averaged for each of the three ROIs and analysed for changes in signal amplitudes and functional connectivity across the relevant speech feedback conditions. For the analysis of signal amplitudes, a general linear model (GLM) was defined that modeled all block types of the respective experiment as separate regressors, which were subsequently delayed and smoothed using a canonical two-gamma hemodynamic response function (HRF) to accommodate for the hemodynamic delay. The GLM was fitted to the region-of-interest data (ROI analysis, see below) of each participant separately (first-level analysis) using ordinary least-squares regression and the ensuing coefficients were subsequently analysed for significant differences at the population level (second-level analysis). To assess the effect of speech feedback in a comparable way across the two studies, we estimated conditional contrasts in the following way. For Study 1, we contrasted overt picture naming without and with noise masking (i.e., PN0 – PNMax), setting all other conditions to 0 (that is, covert PN, Lis0 and LisMax; speech monitoring contrast = [-1 1 0 0 0]). For Study 2, we contrasted overt speaking without noise masking with the maximum noise masking level (i.e., PN0 – PNMax), setting all other conditions to 0, including the intermediate noise levels (i.e., PN1, PN2, Lis0 through LisMax; contrast = [-1 0 0 1 0 0 0 0]).

Functional connectivity between hippocampus and areas involved in speech monitoring were analysed using a psycho- and a physio-physiological interaction model (Friston et al., 1997; O’Reilly et al., 2012). First, we tested whether functional coupling between hippocampus and auditory cortex changed as a function of speech feedback condition (i.e., PN0 > PNMax; *psycho*-physiological interaction, psycho-PI). Here, the contrast between the PN0 and PNMax conditions formed the psychological variable, the mean-centered hippocampal activity as the physiological variable and the point-by-point product between the (mean-centered) psychological and physiological variables as the interaction. Of note, the psychological and interaction terms contained non-zero values only for the timepoints that corresponded to the contrasted conditions, while all other timepoints were set to 0. Second, we tested whether the functional coupling between hippocampus and SMA was correlated with auditory cortex activity during speech feedback conditions (*physio*-physiological interaction, physio-PI (Menon and Levitin, 2005)). Similar to the psycho-PI, the interaction term was obtained by a point-by-point product of the mean-centered physiological variables (i.e., hippocampal and SMA activity). Further, for all PPI variables we set values of the other conditions to 0.

For both PPI models, the (mean-centered) psychological and physiological terms were de-convolved prior to calculating the interaction term, which was then convolved again, using the canonical two-gamma HRF. Finally, to avoid that the other task conditions served as a common source of correlation between the auditory cortex and PPI variables, we appended the task regressors to each of the PPI models.

### Whole-brain physio-PI

In an exploratory analysis, we calculated the whole-brain functional network associated to the hippocampus x SMA connectivity at the voxel-level. To this end, we applied the hippocampus x SMA physio-PI model (of conditions PN0 and PNMax) as described above using a mass-univariate two-level hypothesis test that included all brain voxels. The physio-PI model was appended with task regressors to remove the effects of the task as functional connectivity source. At the first level, functional data were pre-cleaned using the CompCor nuisance terms that we previously estimated. For each participant, the coefficient map of the physio-PI interaction term was smoothed using a Gaussian kernel of 8 mm full-width at half-maximum (FWHM), which then served as entry to the second-level analysis (a mass-univariate voxel-by-voxel one-sample T-test of the subject-level coefficient distribution against 0, df = 22). The second-level statistical map was thresholded at a voxel-level p-value of 0.005 and a minimum cluster size of 1053 mm^3^, as estimated by a cluster threshold algorithm (1000 Monte Carlo simulations using the inherent spatial smoothness of the T-map, at a false-positive rate of 0.05; (Goebel et al., 2006)). Voxel values that survived the statistical thresholds were supervimposed on an anatomical template for visualization.

## Results

### ROI activity

Figure 2 shows the changes in fMRI activity for each of the three ROIs (AC in blue, hippocampus in yellow, SMA in red) when speech feedback was not masked (labeled as *Voice*) and when it was masked with superimposed noise (labeled as *Noise*). For all three ROIs, the distribution of activity values of Study 1 (grey diamonds) overlapped substantially with those of Study 2 (open circles). Statistical analysis confirmed that distributions did not differ between the two studies (two-tailed independent-sample T-tests, df = 21; p(auditory cortex) = 0.12, p(hippocampus) = 0.34, p(SMA) = 0.80). We therefore analysed the combined data of both studies as if they came from the same dataset (aggregated N = 23).

For bilateral auditory cortex, we found significant activity above resting baseline for both speech feedback conditions (PN0 – baseline: T(22) = 5.35, P < 0.001; PNMax – baseline: T(22) = 15.50, P < 0.001). Further, activity decreased when participants heard their own voice as unimpeded speech feedback, compared to when feedback was masked by acoustic noise (PN0 – PNMax: T(22) = -8.28, P < 0.001). These findings replicate the previously published results.

For bilateral hippocampus, we found that activity decreased below resting baseline when participants’ speech feedback was masked (PNMax – baseline: T(22) = - 4.14, P < 0.001), but not when feedback was unimpeded (PN0 – baseline: T(22) = -1.43, P = 0.17). This difference of activity between the two speech feedback conditions was significant (PN0 – PNMax: T(22) = 2.31, P = 0.031). These effects were not previously reported for these datasets, which for Study 1 may have been the result of limited statistical power due to a small sample size in combination with voxel-level multiple comparison corrections (for Study 2, we previously only studied auditory cortical responses).

For bilateral SMA, we found increased activity in each of the two speech feedback conditions (PN0 – baseline: T(22) = 3.74, P = 0.001; PNMax – baseline: T(22) = 3.24, P = 0.004), while there was no significant difference between the two conditions (PN0 – PNMax: T(22) = 0.17, P = 0.87).

When comparing listening conditions with versus without superimposed noise (LIS0 – LISMax), we found no significant differences between the conditions in any of the three ROIs (see Figure 2B; all *ps* > 0.20), suggesting that the feedback effect in auditory cortex and hippocampus only occurred during overt speaking.

### Psycho-physiological interaction (psycho-PI)

Next, we analysed whether the functional coupling between hippocampus and auditory cortex changed as a function of changing speech feedback condition (i.e., PN0 > PNMax). We found that the psycho-physiological interaction term significantly explained auditory cortex activity (controlled for task conditions and hippocampus activity as main effects; T(22) = 2.65, P = 0.015), in which hippocampal-auditory cortex connectivity decreased during noise-masked speech feedback, compared to unimpeded speech feedback. When the psycho-PI was applied to the listening conditions with and without noise masking, the interaction term was not significant (T(22) = -0.95, P = 0.35), indicating that the functional coupling between hippocampus and auditory cortex was specific to overt speech production. Figure 3A schematically displays the psycho-PI effects.

**Figure 3.**
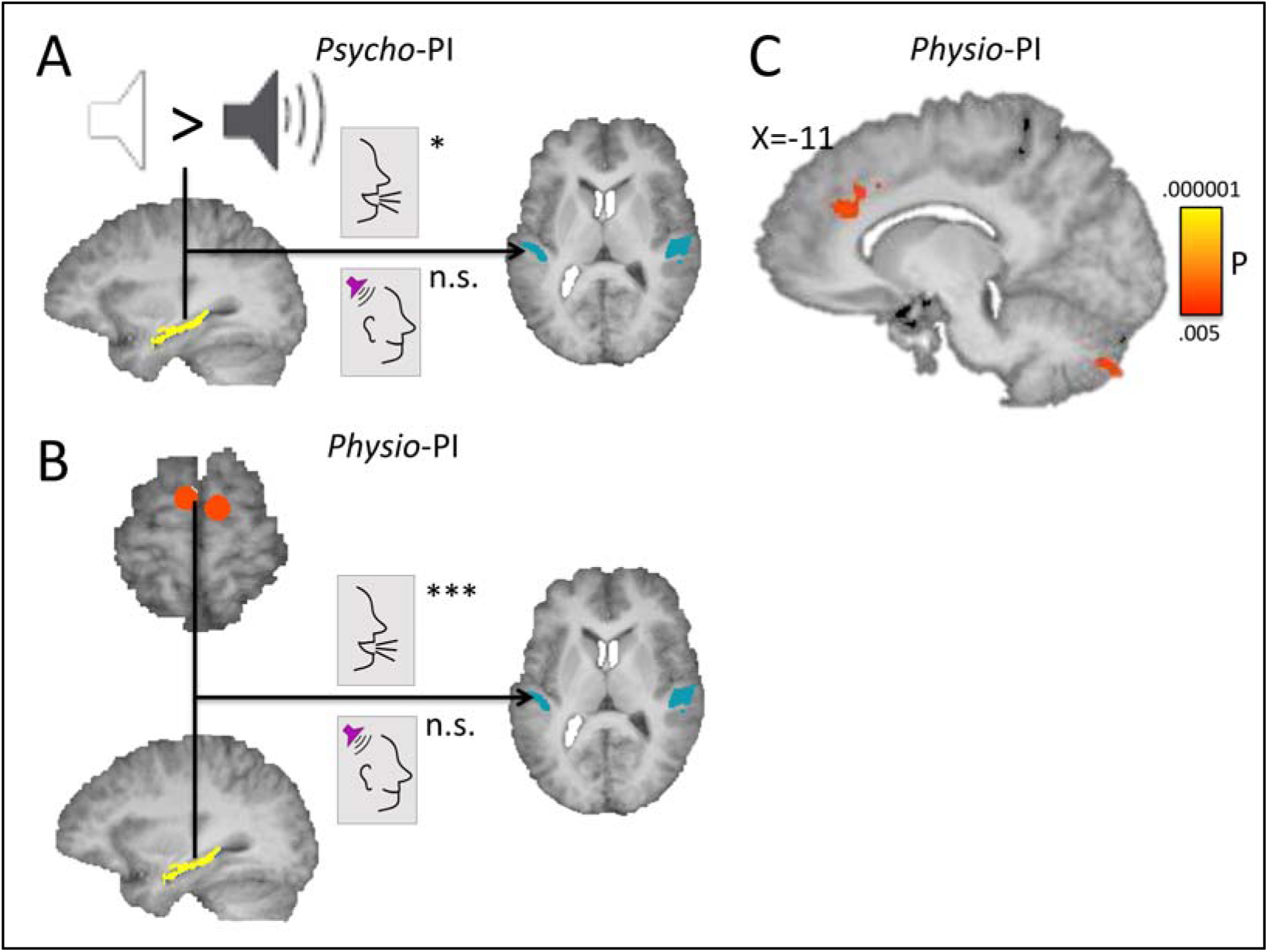
PPI results. A: ROI-based psycho-PI shows significant change in hippocampal-auditory cortex connectivity as a function of speaking condition (PN0 > PNMax) but not during listening. B: ROI-based physio-PI shows significant hippocampal-SMA-auditory functional connectivity during picture naming but not during listening. C: Whole-brain physio-PI shows significant functional connectivity in dorsal ACC and cerebellar cortex. * P < 0.05, *** P < 0.001, n.s. = not significant (P > 0.3).

### Physiological-physiological interaction (physio-PI)

We then analysed whether the functional coupling between SMA and hippocampus explained activity in auditory cortex. We found that the hippocampus x SMA interaction term significantly explained auditory cortex activity during the overt speech conditions (controlled for SMA and hippocampal activity as main effects; T(22) = 4.92, P < 0.001), suggesting that auditory cortex activity was correlated with the functional coupling between SMA and hippocampus during overt speech production (see Figure 3B). When repeating the physio-PI model for the listening blocks with and without noise masking, we found no significant hippocampus x SMA interaction term (T(22) = -0.72, P = 0.48), again suggesting that the functional coupling between hippocampus and SMA was only correlated with auditory cortex activity when participants overtly named the visually presented objects. Thus, merely seeing the objects while not being required to name them did not result in this functional interaction.

To explore the whole-brain functional network of the hippocampus x SMA functional coupling during PN0 and PNMax conditions, we regressed the voxel-level timeseries to the hippocampus x SMA physio-PI model (controlled for main effects of connectivity with hippocampus, SMA and task conditions). Areas that survived statistical thresholding included dorsal anterior cingulate cortex (dACC; x = -15, y = 20, z = 28; size = 1053 mm^3^) and cerebellar cortex (left: -15, -76, -38; 1188 mm^3^; right: 30, - 61, 64; 1107 mm^3^; see Figure 3C). These areas have previously been reported to be involved in speech feedback processing. A physio-PI model that included only listening conditions revealed no significant effects in these areas.

## Discussion

We re-analysed the combined fMRI data collected during overt speech production of two studies and found evidence that human hippocampus is involved in monitoring of speech feedback. Our finding contrasts many older fMRI studies that reported virtually no evidence for a hippocampal involvement in overt picture naming (reviewed in (Indefrey and Levelt, 2000; Price, 2012)), but is in line with more recent evidence for a role of the hippocampus in language processing and verbal communication (Duff et al., 2008; Duff and Brown-Schmidt, 2012; Piai et al., 2016; Kepinska et al., 2018). Further, our observation of larger changes in hippocampal activity when monitoring of speech feedback becomes more challenging fits with other studies that demonstrated that the hippocampus contributes to mismatch processing, showing larger changes in hippocampal activity when encountering novel or unexpected events, compared to expected ones (Kumaran and Maguire, 2006; Long et al., 2016). Thus, we suggest that our results endorse an “expectation”, rather than a “pure memory” account of hippocampal functioning in speech production.

The mismatch process requires that an expectation of future sensory outcomes of speech acts is available. This postulation aligns very well to recent theoretical propositions that hippocampal processing underlies associative prediction of future events or outcomes (Friston and Buzsáki, 2016; Stachenfeld et al., 2017). Further evidence for this suggestion comes from a neurophysiological recording study of hippocampal neurons in human epilepsy patients, which showed increased hippocampal oscillatory activity in the theta range (4-8 Hz) when patients read incrementally presented sentences that provided a strong contextual prediction of the final target word, compared to sentences that provided weak contextual predictions (Piai et al., 2016). In turn, the “expectation” view of hippocampal functioning in speech production can also be applied to interpret previous findings of hippocampal involvement in learning new grammatic rules (Kepinska et al., 2018) or new associations between speech sounds and concepts (Breitenstein et al., 2005), where the novel rules or associations are represented by a mismatch to previously acquired knowledge.

Our finding of decreased hippocampal-auditory cortex connectivity when speech feedback is impaired, compared to unimpeded feedback, also fits to “expectation” view of the hippocampus in speech production. Previous research has shown hippocampal-sensory functional coupling during formation of sensory predictions (Buckner, 2010; Hindy et al., 2016; Kok and Turk-Browne, 2018). In speech production, the increase in auditory cortical activity when speech feedback is impeded or impaired may reflect an increase in error signal resulting from the mismatch between actual and expected sensory feedback (Tourville et al., 2008), but could also reflect a release of inhibition or “neural cancellation”, analogous to the inverse of the mismatch signal (Eliades and Wang, 2005; Christoffels et al., 2011). In both scenarios, the auditory representation during impaired feedback will deviate from the representation of the expected feedback, which could weaken the functional integration with other brain areas that also contribute to the processing of that information (Obleser et al., 2007; Nath and Beauchamp, 2011). The decrease in functional auditory-hippocampal coupling that we observed could reflect the increased discordance between auditory and hippocampal representations of sensory expectations during speaking.

Further support for a hippocampal involvement comes from our finding that auditory cortical activity could be explained by a hippocampal-SMA interaction during speech production, but not during listening to one’s own pre-recorded voice. Whole-brain analysis further showed that the hippocampal-SMA interaction explained activity in dorsal ACC and cerebellar cortex (beyond connectivity of these areas with either hippocampus or SMA). Ample research has implicated SMA in speech control (Abel et al., 2009; Tremblay and Gracco, 2009; van de Ven et al., 2009; Gauvin et al., 2016; Botha et al., 2018), where speech monitoring may arise from the functional coupling between SMA and auditory cortex. Particularly, connectivity may increase when speech feedback is masked (van de Ven et al., 2009), while lower connectivity may be related to more articulatory errors in speech disorders (Botha et al., 2018). The SMA is a candidate region to provide a forward model of motor acts that is used to predict sensory consequences (Wolpert et al., 1995). Our finding suggests that the hippocampus may contribute to mismatch processing by providing an associative mismatch signal that incorporates information about the predicted and actual sensory outcomes in varying acoustic contexts. Changes in hippocampal activity or connectivity could result from integrating the sensory prediction derived from the forward model with the perceived sensory consequences, which in turn may be used to update or augment the speech production process. The whole-brain connectivity results, finally, suggest that a mismatch signal that is coded by the hippocampal-SMA interaction may be further processed by ACC and cerebellum, which have previously been reported to be involved in speech production tasks (Hirano et al., 1997; Christoffels et al., 2007; Abel et al., 2009). The ACC has been associated to error monitoring (Van Veen and Carter, 2002), which suggests that, during speaking, it may process the mismatch signal that is encoded within the hippocampal-SMA coupling. The role of the cerebellum in multi-sensory (Naumer et al., 2010) and sensori-motor convergence (Paulin, 1993; Brooks and Cullen, 2019) suggests that it contributes to the integration of speech feedback with expected sensory consequences of motor acts. Finally, connectivity between hippocampus and ACC has been related to mapping of context-dependent action-outcome representations (Rolls, 2019). In all, these findings suggest that processing of mismatch between expected and actual sensory consequences of speaking entail interaction between motor and associative representations of speech production and auditory sensory representations of perceived speech feedback, while the degree of mismatch may be further processed in higher-order areas.

It should be noted here that participants made very few errors (less than 1% of the trials), which limits our inferences about the role of the hippocampus in speech monitoring. For example, it is possible that the hippocampal contribution to speech monitoring allowed task performance to remain very high, but experimental testing of this proposition would require a higher error rate. Further, while we endorse an “expectation” view of the hippocampus in speech production, we do not exclude the contribution of memory (Klooster and Duff, 2015). However, the key issue in our argumentation is that a “pure memory” account falls short of explaining changes in hippocampal activity when speech feedback is impaired. Future studies are needed to further disentangle semantic memory from associative mismatch processing in speech production.

In conclusion, we showed that the hippocampus is involved in the online monitoring the sensory consequences of our own speech acts. Our findings are in line with recent propositions that the hippocampus plays a role in language production and verbal communication (Duff et al., 2008; Duff and Brown-Schmidt, 2012; Friston and Buzsáki, 2016; Piai et al., 2016), and show that the hippocampus may be involved in predictive processing of speech production, rather than or beyond contributing semantic memory representations of what is said.

